# *De novo* Mutations in *NALCN* Cause a Syndrome of Congenital Contractures of the Limbs and Face with Hypotonia, and Developmental Delay

**DOI:** 10.1101/013656

**Authors:** Jessica X. Chong, Margaret J. McMillin, Kathryn M. Shively, Anita E. Beck, Colby T. Marvin, Jose R. Armenteros, Kati J. Buckingham, Naomi T. Nkinsi, Evan A. Boyle, Margaret N. Berry, Maureen Bocian, Nicola Foulds, Maria Luisa Giovannucci Uzielli, Chad Haldeman-Englert, Raoul C.M. Hennekam, Paige Kaplan, Antonie D. Kline, Catherine L. Mercer, Malgorzata J.M. Nowaczyk, Jolien S. Klein Wassink-Ruiter, Elizabeth W. McPherson, Regina A. Moreno, Angela E. Scheuerle, Vandana Shashi, Cathy A. Stevens, John C. Carey, Arnaud Monteil, Philippe Lory, Holly K. Tabor, Joshua D. Smith, Jay Shendure, Deborah A. Nickerson, Michael J. Bamshad

**Author notes:** Corresponding author: Mike Bamshad, MD, Department of Pediatrics, University of Washington School of Medicine, Box 356320, 1959 NE Pacific Street, HSB RR349, Seattle, WA 98195.

## Abstract

Freeman-Sheldon syndrome, or distal arthrogryposis type 2A (DA2A), is an autosomal dominant condition caused by mutations in *MYH3* and characterized by multiple congenital contractures of the face and limbs and normal cognitive development. We identified a subset of five simplex cases putatively diagnosed with “DA2A with severe neurological abnormalities” in which the proband had Congenital Contractures of the LImbs and FAce, Hypotonia, and global Developmental Delay often resulting in early death, a unique condition that we now refer to as CLIFAHDD syndrome. Exome sequencing identified missense mutations in *sodium leak channel, nonselective* (*NALCN*) in four families with CLIFAHDD syndrome. Using molecular inversion probes to screen *NALCN* in a cohort of 202 DA cases as well as concurrent exome sequencing of six other DA cases revealed *NALCN* mutations in ten additional families with “atypical” forms of DA. All fourteen mutations were missense variants predicted to alter amino acid residues in or near the S5 and S6 pore-forming segments of NALCN, highlighting the functional importance of these segments. *In vitro* functional studies demonstrated that mutant NALCN nearly abolished the expression of wildtype NALCN, suggesting that mutations that cause CLIFAHDD syndrome have a dominant negative effect. In contrast, homozygosity for mutations in other regions of *NALCN* has been reported in three families with an autosomal recessive condition characterized mainly by hypotonia and severe intellectual disability. Accordingly, mutations in *NALCN* can cause either a recessive or dominant condition with varied though overlapping phenotypic features perhaps depending on the type of mutation and affected protein domain(s).

---

Distal arthrogryposis (DA) is a group of at least ten disorders characterized by non-progressive, congenital contractures of two or more body areas that typically affect the wrists, hands, ankles, and feet^1^. The three most common DA syndromes are DA1 (MIM 108120)^1,2^, DA2A (Freeman-Sheldon syndrome; MIM 193700)^3^, and DA2B^1,2^, (Sheldon-Hall syndrome; MIM 601680). DA1 and DA2B can be caused by variants in any one of several genes including *TPM2* (MIM 190990), *TNNT3* (MIM 600692), *TNNI2* (MIM 191043), and *MYH3* (MIM 160720),^4^ whereas DA2A is only caused by mutations in *MYH3.* Moreover, mutations in *MYH3* are found in more than ninety percent of persons who meet the diagnostic criteria for DA2A^5^.

Among persons with DA2A referred to our research program over the past decade in whom no pathogenic mutations were identified in *MYH3,* we recognized a pattern of clinical characteristics among a subset of individuals suggestive of a distinctive and previously unrecognized multiple malformation syndrome. Specifically, we identified a subset of five families with an affected child born to unaffected parents in which the proband had Congenital Contractures of the LImbs (hands, feet) and FAce; abnormal tone most commonly manifest as Hypotonia; neonatal respiratory distress; and Developmental Delay, or CLIFAHDD syndrome (Table 1, Families A, B, C, D, and E; Figure 1, Figures S1 and S2).

**Table 1.** Mutations and Clinical Findings of Individuals with CLIFAHDD Syndrome. This table provides a summary of clinical features of affected individuals from families in which mutations in *NALCN* were identified. Clinical characteristics listed in table are primarily features that distinguish CLIFAHDD syndrome from Distal Arthrogryposis conditions. Plus (+) indicates presence of a finding, minus (-) indicates absence of a finding, ND = no data were available. N/A = not applicable. GERP = Genomic Evolutionary Rate Profiling. CADD = Combined Annotation Dependent Depletion. CLIFAHDD = <u>C</u>ongenital contractures of the <u>LI</u>mbs, <u>FA</u>ce, <u>H</u>ypotonia and <u>D</u>evelopmental Delay. DA2A = Distal arthrogryposis type 2A. DA2B = Distal arthrogryposis type 2B. DA1 = Distal arthrogryposis type 1. VSD = ventricular septal defect. GERD = Gastroesophageal reflux disease. cDNA positions provided as named by the HGVS MutNomen web tool relative to NM_052867.2

| Family | A | B | C | D | E | F | G |
| --- | --- | --- | --- | --- | --- | --- | --- |
| **Original Diagnosis** | CLIFAHDD | CLIFAHDD | CLIFAHDD | CLIFAHDD | CLIFAHDD | DA2A | DA2A |
| **Mutation Information** |  |  |  |  |  |  |  |
| Exon (NALCN) | 6 | 9 | 15 | 39 | 31 | 8 | 13 |
| cDNA change | c.530A>C | c.938T>G | c.1768C>T | c.4338T>G | c.3493A>C | c.934C>A | c.1526T>C |
| Predicted protein alteration | p.Gln177Pro | p.Val313Gly | p.Leu590Phe | Ile1446Met | p.Thr1165Pro | p.Leu312Ile | p.Leu509Ser |
| GERP | 4.92 | 6.16 | 5.94 | 4.73 | 5.2 | 6.16 | 5.11 |
| CADD 1.0 (phred-like) | 19.22 | 25.9 | 23.9 | 16.08 | 21.0 | 32.0 | 22.6 |
| Polyphen-2 (HumVar) | 1.0 | 1.0 | 0.996 | 0.995 | 0.969 | 1.0 | 1.0 |
| **Clinical Features** |  |  |  |  |  |  |  |
| **Face** |  |  |  |  |  |  |  |
| Downslanting palpebral fissures | — | + | ND | + | + | + | + |
| Strabismus | — | + | ND | + | + | + | + |
| Esotropia | — | + | ND | — | + | — | — |
| Broad nasal bridge | + | + | + | + | + | + | + |
| Anteverted nasal tip | + | + | + | + | + | — | + |
| Large nares | + | + | + | + | + | + | + |
| Short columella | + | + | + | + | + | + | + |
| Long philtrum | + | + | + | + | + | + | + |
| Micrognathia | + | + | + | + | + | + | + |
| Pursed lips | + | + | + | + | + | — | + |
| H-shaped dimpled chin | + | + | + | + | — | + | + |
| Deep nasolabial folds | + | + | + | + | + | — | + |
| Full cheeks | + | + | + | + | + | + | — |
| **Limbs** |  |  |  |  |  |  |  |
| Camptodactyly | + | + | + | + | + | + | + |
| Ulnar deviation | + | + | + | + | + | + | + |
| Adducted thumbs | + | + | + | + | + | + | + |
| Clubfoot | + | + | + | L | Metatarsus adductus | + | + |
| Calcaneovalgus deformity | — | + | — | R | — | — | — |
| Hip contractures | + | + | + | BL dislocation | — | — | + |
| Elbow contractures | + | + | + | — | — | + |  |
| Knee contractures | + | — | + | + | — | + | + |
| **Other** |  |  |  |  |  |  |  |
| Scoliosis | + | + | — | + | — | ND | + |
| Short neck | + | + | + | + | + | ND | + |
| Cognitive delay | ND* | + | ND* | + | + | + | – |
| Speech delay | N/A | + | N/A | + | + | + | + |
| Motor delay | + | + | + | + | + | + | + |
| Seizures | + | + | – | – | – | – | – |
| Questionable seizure activity or abnormal breathing, loss motor control | ND | + | ND | abnormal discharges | periods of rigidity | + | + |
| MRI findings | Normal | Mild cerebral and cerebellar atrophy | ND | Generalized cerebral atrophy, small pituitary | Normal | Cerebellar atrophy | normal |
| Hypotonia | no | yes | yes | yes | yes | yes | no |
| Respiratory insufficiency | + | + | + | + | + | apneic episodes in newborn period | – |
| Abnormal respiratory pattern in newborn period | + | + | + | + | + | + | – |
| Excessive drooling | ND | ND | ND |  |  | + |  |
| GERD | ND | + | ND | + | + | + | ND |
| Constipation | ND | occasional | ND | – |  | + |  |
| Hernia | – | Inguinal | Inguinal | Inguinal / peri umbilical | Inguinal | – | Inguinal |
| Age at time of death | 6 months | 5 years | 4 months | N/A | N/A | N/A | N/A |
| other |  | Central apnea, uric acid renal stones | 11 pairs of ribs, abnormal pelvis and scapulae | Renal calcifications |  | Suspected fatty acid oxidation defect |  |

| Family | H | I | J | K | L | M | N |
| --- | --- | --- | --- | --- | --- | --- | --- |
| **Original Diagnosis** | DA2A | DA2B | DA2B | DA2B | DA2B | DA1 | DA1 |
| **Mutation Information** |  |  |  |  |  |  |  |
| Exon (NALCN) | 26 | 9 | 13 | 14 | 31 | 13 | 26 |
| cDNA change | c.3017T>C | c.979G>A | c.1538C>A | c.1733A>C | c.3542G>A | c.1534T>G | c.3050T>C |
| Predicted protein alteration | p.Val1006Ala | p.Glu327Lys | p.Thr513Asn | p.Tyr578Ser | p.Arg1181Gln | p.Phe512Val | p.Ile1017Thr |
| GERP | 5.03 | 6.16 | 5.11 | 5.3 | 5.2 | 5.11 | 5.03 |
| CADD (phred-like) | 24.2 | 36.0 | 25.5 | 23.7 | 20.5 | 24.6 | 17.54 |
| Polyphen-2 (HumVar) | 0.999 | 1.0 | 1.0 | 1.0 | 0.444 | 1.0 | 1.0 |
| **Clinical Features** |  |  |  |  |  |  |  |
| **Face** |  |  |  |  |  |  |  |
| Downslanting palpebral fissures | + | ? | + | + | + | – | + |
| Strabismus | + | – | + | – | – | – | – |
| Esotropia | – | – | – | + | – | – | – |
| Broad nasal bridge | + | + | + | + | + | + | + |
| Anteverted nasal tip | + | + | + | – | + | + | + |
| Large nares | + | + | + | + | + | + | + |
| Short columella | + | + | + | + | + | + | + |
| Long philtrum | + | + | + | + | – | – | + |
| Micrognathia | + | + | + | + | + | + | – |
| Pursed lips | – | – | + | + | + | – | ? |
| H-shaped dimpled chin | – | – | + | – | + | – | – |
| Deep nasolabial folds | + | + | – | + | + | + | + |
| Full cheeks | + | + | + | + | + | + | + |
| **Contractures** |  |  |  |  |  |  |  |
| Camptodactyly | + | + | + | + | + | + | + |
| Ulnar deviation | + | + | + | + | + | + | + |
| Adducted thumbs | + | + | + | + | + | + | + |
| Clubfoot | Mild L varus | + | + | + | – | Mild varus | + |
| Calcaneovalgus deformity | – | – | – | – | – | + | – |
| Hip contractures | – | + | + | + | + | Internally rotated | + |
| Elbow contractures | + | + | – |  | + | – | – |
| Knee contractures | + | + | – | + | + | – | – |
| **Other** |  |  |  |  |  |  |  |
| Scoliosis | – | – | – | + | – | Lumbar | – |
|  |  |  |  |  |  | lordosis |  |
| Short neck | – | + | – | + | + | + |  |
| Cognitive delay | + | + | + | + | + | + | + |
| Speech delay | + | + | + | + | + | + | + |
| Motor delay | + | + | + | + | + | + | + |
| Seizures | – | – | – | – | – | – | – |
| Questionable seizure activity or abnormal breathing, loss motor control | ND | ND | + | + | Seizure-like activity with fever | ND | Irregular breathing |
| MRI findings | ND | ND | Diffuse cerebellar atrophy | ND | normal | normal | White matter edema |
| Hypotonia | yes | ND | no | no | no? | yes | no |
| Respiratory insufficiency | ND | ND | ND | + | ND | – | + |
| Abnormal respiratory pattern in newborn period | ND | ND | ND | + | + | – | + |
| Excessive drooling | ND | ND | + | + | + | ND | ND |
| GERD | – | + | + | + | + | In infancy | – |
| Constipation | ND | ND | + | + | + | ND | ND |
| Hernia | – | Inguinal | Inguinal | Inguinal | ND | Umbilical | – |
| Age at time of death | N/A | N/A | N/A | N/A | N/A | N/A | N/A |
| other |  | Cleft palate |  | Mitral valve prolapse |  | Small VSD |  |

Child died before assessment could be made

**Figure 1.**
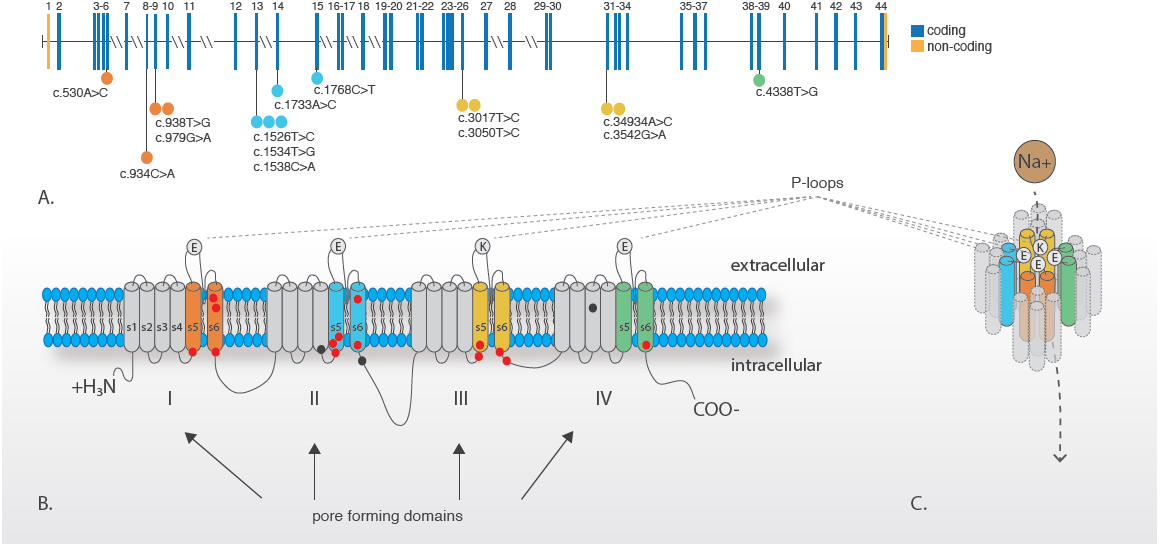
Phenotypic Characteristics of Each Individual with CLIFAHDD Syndrome. Four individuals affected with CLIFAHDD syndrome (A-I); all individuals shown have *NALCN* mutations. Note the short palpebral fissures, flattened nasal root and bridge, large nares, long philtrum, pursed lips, H-shaped dimpling of the chin and deep nasolabial folds (A, B, C-1, D). Camptodactyly of the digits of the hands and ulnar deviation of the wrist (A, B, C, D-1, D-2) or clubfoot (C-3) is present in each affected person. Case identifiers for the individuals shown in this figure correspond to those in Table 1, where there a detailed description of the phenotype of each affected individual. Figure S1 provides a pedigree of each CLIFAHDD family.

Facial characteristics shared among individuals with CLIFAHDD syndrome included downslanting palpebral fissures, broad nasal bridge with anteverted nasal tip and large nares, a short columella, long philtrum, deep nasolabial folds, micrognathia, pursed lips, and dimpling of the chin resembling the “H-shaped” chin observed in persons with DA2A (Figure 1). Each person had camptodactyly with adducted thumbs, as well as positional foot deformities ranging from mild varus deformity to severe clubfoot. Additionally, contractures of the elbows, knees, and hips, as well as short neck and scoliosis were observed.

Neurological evaluation of each living child with CLIFAHDD revealed global developmental delay manifest as speech, motor, and cognitive delays that varied from mild to severe, as well as hypotonia and seizures. Magnetic resonance imaging of the brain in four of the affected individuals was reported as normal in two persons and abnormal in two. Of the latter, one person was reported to have cerebral and cerebellar atrophy and another to have cerebral atrophy and a “small pituitary.” Severe gastroesophageal reflux was observed in all three individuals for whom data were available, and four of five individuals had inguinal hernias. Three of the five affected individuals died in infancy or early childhood. One person died of complications of hemophagocytosis in the newborn period while the exact cause of death was unclear in the other two individuals.

To find the gene(s) for CLIFAHDD syndrome, we first screened the proband of each of the five families putatively diagnosed with CLIFAHDD syndrome by Sanger sequencing for mutations in genes other than *MYH3* known to cause DA1, DA2A and DA2B. Each proband with CLIFAHDD syndrome was also screened for copy number variations (CNVs) by comparative genome-wide array genomic hybridization on the Illumina HumanCytoSNP-12. No pathogenic mutations or shared CNVs were identified. Next, exome sequencing was performed on four CLIFAHDD parent-child trios for whom sufficient quantities of DNA were available. All studies were approved by the institutional review boards of the University of Washington and Seattle Children’s Hospital, and informed consent was obtained from participants or their parents.

One microgram of genomic DNA was subjected to a series of shotgun library construction steps, including fragmentation through acoustic sonication (Covaris), end-polishing (NEBNext End Repair Module), A-tailing (NEBNext dA-Tailing Module), and PCR amplification with ligation of 8 bp barcoded sequencing adaptors (Enzymatics Ultrapure T4 Ligase) for multiplexing. One microgram of barcoded shotgun library was hybridized to capture probes targeting ~36.5 Mb of coding exons (Roche Nimblegen SeqCap EZ Human Exome Library v2.0) Library quality was determined by examining molecular weight distribution and sample concentration (Agilent Bioanalyzer). Pooled, barcoded libraries were sequenced via paired-end 50 bp reads with an 8 bp barcode read on Illumina HiSeq sequencers.

Demultiplexed BAM files were aligned to a human reference (hg19) using the Burrows-Wheeler Aligner (BWA) 0.6.2. Read data from a flow-cell lane were treated independently for alignment and QC purposes in instances where the merging of data from multiple lanes was required. All aligned read data were subjected to: (1) removal of duplicate reads (Picard MarkDuplicates v1.70) (2) indel realignment (GATK IndelRealigner v1.6-11-g3b2fab9); and (3) base quality recalibration (GATK TableRecalibration v1.6-11-g3b2fab9). Variant detection and genotyping were performed using GATK UnifiedGenotyper (v1.6-11-g3b2fab9). Variant data for each sample were formatted (variant call format [VCF]) as “raw” calls that contained individual genotype data for one or multiple samples, and flagged using the filtration walker (GATK) to mark sites that were of lower quality and potential false positives (e.g. strand bias > −0.1, quality scores (Q50), allelic imbalance (ABHet≥0.75), long homopolymer runs (HRun>3), and/or low quality by depth (QD<5).

Variants with an alternative allele frequency >0.005 in the ESP6500 or 1000 Genomes or >0.05 in an internal exome database of ~700 individuals were excluded prior to analysis. Additionally, variants that were flagged as low quality or potential false positives (quality score ≤ 30, long homopolymer run > 5, low quality by depth < 5, within a cluster of SNPs) were also excluded from analysis. Variants that were only flagged by the strand bias filter flag (strand bias > −0.10) were included in further analyses as the strand bias flag has previously been found to be applied to valid variants. CNV calls were also generated from exome data with CoNIFER^6^. Variants were annotated with the SeattleSeq137 Annotation Server, and variants for which the only functional prediction label was “intergenic,” “coding-synonymous,” “utr,” “near-gene,” or “intron” were excluded. Individual genotypes with depth<6 or genotype quality<20 were treated as missing in analysis.

Analysis of variants from exome sequencing under a *de novo* mutation model identified four different *de novo* variants in a single gene, *NALCN,* encoding a sodium leak channel (NALCN [MIM 611549; Refseq NM_052867.2]) in four families (Table 1 and Figure 1, Families A, B, C, and D; Figures S1 and S2) with CLIFAHDD syndrome. Concurrent exome sequencing of six trios with DA in whom no pathogenic variants had been identified previously in genes known to cause DA serendipitously revealed *de novo* variants in *NALCN* in two families (Table 1, Families G and K; Figure S3). All six variants were missense variants predicted to be deleterious and result in amino acid substitutions of highly conserved amino acid residues (i.e., the minimum Genomic Evolutionary Rate Profile score was 4.73) in *NALCN* (Table 1 and Figure 2). Each variant was confirmed by Sanger sequencing to have arisen *de novo* and none of the six variants were found in over 71,000 control exomes comprised of the ESP6500, 1000 Genomes phase 1 (Nov 2010 release), internal databases (>1,400 chromosomes), or ExAC (October 20, 2014 release).

**Figure 2.**
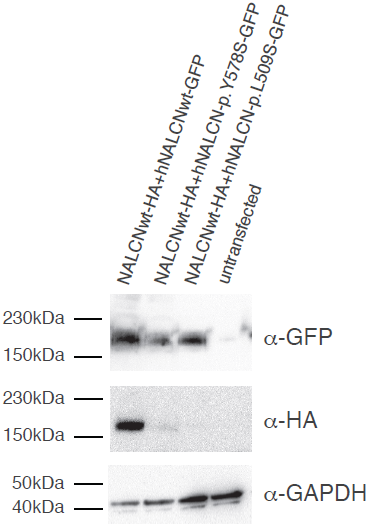
Genomic and protein structure of NALCN and Spectrum of mutations that cause CLIFAHDD syndrome. A) *NALCN* is composed of 45 exons, 43 of which are protein-coding (blue) and two of which are non-coding (orange). Lines with attached dots indicate the approximate locations of fourteen different *de novo* variants that cause CLIFAHDD syndrome. The color of each dot reflects the nearest transmembrane segment (S1-S6) depicted in B and C. The two mutations between IIIS6 and IVS1 are located in the intracellular loop linking domains III and IV, which is known to be involved in the inactivation of voltage-gated sodium channels; however NALCN is not voltage-gated, so the specific function of this loop in NALCN is not known. B) Predicted protein topology of NALCN is comprised of four homologous pore-forming domains (I—IV) each composed of six transmembrane segments (S1-S6). Bold line represents the polypeptide chain linking segments and domains. The outer ring of amino acid residues of each P-loop that form the ion selectivity filter (EEKE) is represented by filled circles. The approximate positions of variants that cause CLIFAHDD syndrome and infantile hypotonia with psychomotor retardation and characteristic facies are indicated by red filled circles and black filled circles, respectively. C) S1-S3 may have structural functions, while four pore-forming loops (P-loops) spanning from S5 to S6 form the ion selectivity filter. In other 4x6TM protein superfamily channels, such as SCN1A, the S4 segments initiate opening of the central pore (S5/Ploop/S6) in response to voltage changes, however because NALCN is not voltage-gated, the function of S4 is currently unknown.

Because the phenotype of the individuals with mutations in *NALCN* was variable and overlapped with other DA syndromes, we used molecular inversion probes with 5-bp molecular tags^7^ (smMIPs; designed with MIPGEN^8^ v.0.9.7) to conduct targeted next-generation sequencing of *NALCN* in 202 additional samples from persons with DA in which no pathogenic mutation had been found. The 43 coding exons (5,214 bp total) of Ensembl transcript ENST00000251127 and 10 bp flanking each exon (860 bp total) were targeted using smMIPs, for an overall target size of 6,074 bp. Pooled and phosphorylated smMIPs were added to the capture reactions with 100 ng of genomic DNA from each individual to produce a *NALCN* library for each individual. The libraries were amplified during 21 cycles of PCR, during which an 8 bp sample barcode was introduced. The barcoded libraries were then pooled and purified using magnetic beads. After Picogreen quantification to determine the appropriate dilution, 10 picomoles of the pool was sequenced on an Illumina MiSeq. MIP collapsing and arm trimming (MIPGEN v1.0), alignment (BWA 0.7.8), and multi-sample genotype calling (GATK Unified Genotyper 3.2-2-gec30cee) were performed and variants were annotated with SeattleSeq138. We used the same filtering strategy employed in our analysis of the exome sequences to select variants for further confirmation by Sanger sequencing.

Missense variants predicted to result in substitution of conserved amino acid residues were identified and confirmed to have arisen *de novo* in eight additional families—one family with CLIFAHDD syndrome that had not been available for exome sequencing (Table 1), two families originally diagnosed with DA2A (Table 1, Families F and H; Figure S1; Figure S3, Family F), three families originally diagnosed with DA2B (Table 1, Families I, J and L; Figure S1; Figure S2, Families I and J), and two families diagnosed with DA1 (Table 1, Families M and N; Figure S1; Figure S3, Family M). Collectively, we discovered unique, *de novo* missense mutations in *NALCN* in fourteen simplex families in which the proband had been diagnosed with either CLIFAHDD syndrome (n=5), DA2A (n=3), DA2B (n=4), or DA1 (n=2) (Table 1 and Figure 2).

The diagnosis of CLIFAHDD syndrome was considered in one additional case of DA2A referred to our center based on the presence of a pattern of congenital facial and limb contractures characteristic of DA2A in the absence of a finding of a pathogenic *MYH3* mutation. However, death occurred within two hours of birth at 29 weeks gestation, and, therefore, the available clinical information was considered too limited. This approach proved prudent as no mutation in *NALCN* was identified by exome sequencing. Instead, this child was found to be a compound heterozygote in *RYR1* (MIM 180901) for a predicted nonsense variant, p.(Tyr3921Ter) (NM_000540.2:c.11763C>A) inherited from the father, and a missense variant, p.(Gly341Arg) (NM_000540.2:c.1021G>A), inherited from the mother. The missense variant has been previously reported to cause malignant hyperthermia, and persons carrying this variant had a positive “in vitro contractility test.”^9^ Crisponi syndrome^10,11^ (MIM 601378) was also considered as a possible diagnosis for this child. However no candidate variants were identified in *CRLF1* (MIM 604237) in this child or any of the fourteen families in which *NALCN* mutations were identified. Last, we have previously found two mutations in *RYR1* (i.e., c.1391A>G, p.Gln464Arg and c.2683-1G>A) in a child diagnosed with DA2A, suggesting that mutations in *RYR1* are a rare cause of DA2A.

Mutations in *NALCN* have recently been reported to cause an autosomal recessive condition, infantile hypotonia with psychomotor retardation and characteristic facies (IHPRF; MIM 615419), in one Turkish^12^ and two Saudi Arabian^13^ families. Individuals affected with CLIFAHDD syndrome share some of the phenotypic characteristics of IHPRF (e.g., developmental delay and hypotonia). However, these are relatively non-specific findings, and IHPRF and CLIFAHDD syndrome appear to be otherwise distinct from one another. In addition, the parents of children with IHPRF are carriers of *NALCN* mutations, but reportedly do not have any of the abnormalities found in persons with CLIFAHDD syndrome. Furthermore, each inherited variant found in *NALCN* in the fourteen individuals with CLIFAHDD syndrome was also present in either ESP6500 or 1000 Genomes data. This finding suggests that these inherited variants are polymorphisms rather than pathogenic variants underlying CLIFAHDD syndrome. Together, these results support the hypothesis that CLIFAHDD syndrome is an autosomal dominant condition distinct from IHPRF.

We next sought to explore the mechanism by which variants in *NALCN* result in CLIFAHDD syndrome. To this end, we constructed two expression plasmids carrying *NALCN* variants: pcDNA3-hNALCN-L509S-GFP with p.(Leu509Ser) found in a person with relatively mild developmental delay (Table 1, Family G) and pcDNA3-hNALCN-Y578S-GFP with p.(Tyr578Ser) found in a child with severe developmental delay (Table 1, Family K) and co-expressed mutant and wild-type *NALCN* in HEK-293T cells. Western blot analysis demonstrated that the expression of either p.(Tyr578Ser) or p.(Leu509Ser) (GFP-tagged) *NALCN* mutants nearly eliminated wild-type (HA-tagged) NALCN protein (Figure 3).

**Figure 3.** Expression of p.(Tyr578Ser) and p.(Leu509Ser) NALCN mutants prevent wildtype NALCN detection at the protein level when co-expressed in HEK-293T cells. HA-tagged wild-type NALCN channels are not detectable when co-expressed with GFP-tagged NALCN mutant channels in HEK-293T cells but are detectable when co-expressed with GFP-tagged wild-type NALCN channels. This experiment is representative of four independent experiments.

It remains to be determined whether synthesis of mutated NALCN channels prevents wild-type NALCN synthesis or induces its degradation by the proteasome. The latter mechanism has been previously described^14^ for mutations in the Cav1.2 calcium channel responsible for dominantly-inherited Episodic Ataxia type 2 (MIM 108500). However, preliminary evidence suggests that NALCN mutants are correctly targeted to the plasma membrane (data not shown). Nevertheless, both mutations induced disappearance of wild-type NALCN to apparently the same extent. Accordingly, an explanation for the phenotypic differences observed between the cases sampled is not yet apparent.

Collectively, these functional studies, while preliminary, are consistent with mutations in *NALCN* causing CLIFAHDD syndrome via a dominant negative effect that results in haploinsufficiency. However, the finding that persons heterozygous for *NALCN* mutations that cause IHPRF are reportedly unaffected indicates that haploinsufficiency of NALCN alone is not adequate to cause CLIFAHDD syndrome. These observations suggest that *NALCN* mutations that cause CLIFAHDD syndrome may produce a mutant protein that has some residual activity (i.e. a hypomorphic allele) or that exhibits a gain of function. Moreover, mutations in *NALCN* could cause CLIFAHDD syndrome by more than one mechanism.

NALCN is a G protein-coupled receptor-activated channel^15^ consisting of four homologous domains (i.e., domains I-IV), each of which consists of six transmembrane segments (S1-S6), separated by cytoplasmic linkers (Figure 2). Pore-forming loops (P-loops) between S5 and S6 of each domain form an EEKE sodium ion selectivity filter (Figure 2). In vertebrates and some invertebrate model organisms such as *Drosophila* and *C. elegans,* NALCN exclusively has a sodium-selective EEKE pore and putatively functions as a sodium channel. However, it remains unclear whether NALCN is actually an ion channel rather than a sensor of sodium^16^. In mammals, NALCN is most highly expressed in the central nervous system but is also found at moderate levels in heart, lymph nodes, pancreas, and thyroid (summarized by ^17^). Unlike most known genes underlying DA syndromes, *NALCN* is not expressed in fetal skeletal muscle (Figure S4). However, homologues of *NALCN* are expressed in the neuromuscular junction in *D. melanogaster*^18^ as well as in motor neurons in *C. elegans*^19^ and mice^20,21^. Accordingly, mutations in NALCN may cause congenital contractures by disturbing motor control of myofiber function during development similar to the hypothesized effects of mutations in *ECEL1, PIEZO2* and *CHRNG* (MIM 100730).

Each of the fourteen *de novo* mutations in *NALCN* found to cause CLIFAHDD syndrome is located in or near the predicted S5 and S6 segments that are part of the pore forming domain of the NALCN channel (Figure 2; Table S1). The clustering of mutations is an indication of the functional importance of these segments. In contrast, the frameshift (c.1489delT [p.Tyr497Thrfs21]) and nonsense (c.1924C>T [p.Gln642Ter]) mutations reported to cause IHPRF are predicted to result in loss of function of *NALCN* while the missense (c.G3860T [p.Trp1287Leu]) mutation is predicted to alter the S3 segment of domain IV. These findings suggest that mutations in different regions of *NALCN* perturb different functions of the channel, result in different Mendelian conditions, and determine whether the condition is transmitted in an autosomal recessive versus an autosomal dominant pattern. A similar relationship has recently been reported for *MAB21L2* (MIM 604357) mutations that underlie a spectrum of major malformations of the eye.^22^

NALCN is highly conserved across vertebrates, with ~98% sequence identity between human and mouse NALCN, and functional studies in model organisms have illuminated multiple physiologic roles of NALCN. In general, NALCN permits background sodium ion leak current in neurons that contribute to regulation of resting membrane potentials and spontaneous firing of motor neurons^23^. In mice, NALCN has been shown to regulate respiration^24^, intestinal pacemaking activity in the interstitial cells of Cajal^25^, and serum sodium concentration^26^. Mutation of *NALCN* homologs in *D. melanogaster, C. elegans,* and *L. stagnalis* results in abnormal locomotion^19,27-30^, disturbance of circadian rhythms^29^, disruption of normal respiratory rhythms^31^, and altered sensitivity to ethanol and some general anesthetics (^30,32,33^; reviewed by ^17^). Homozygous *Nalcn* null mice die shortly after birth and exhibit disrupted respiratory rhythm, with mutant pups breathing for five seconds alternating with five seconds of apnea, as a result of a lack of electrical discharges from the C4 nerves that control the thoracic diaphragm^24^. Interstitial cells of Cajal, which generate the electrical signals that govern gut motility, from *Nalcn* null mice exhibit abnormal regulation of pacemaking activity^25^. These abnormalities are reminiscent of some of the respiratory and gastrointestinal abnormalities observed in persons with CLIFAHDD syndrome. Moreover, gain of function mutations in exons encoding segment S6 of domain I of a *NALCN* homolog (*nca-1*) in *C. elegans* lead to a dominantly-inherited phenotype that includes pausing and uncoordinated, exaggerated body bending^19,34^ and loosely recapitulates episodes of disorganized movement experienced by several individuals with CLIFAHDD syndrome. Nevertheless, these assertions are based on anecdotal clinical information and necessitate objective testing of respiratory function and gastrointestinal motility.

NALCN is part of a structurally-similar superfamily of cation (i.e., sodium and calcium) channels with four homologous domains containing six transmembrane helices, the 4x6TM family^16^. Mutations in 4x6TM genes that encode voltage-gated sodium and calcium channels (e.g., *SCN1A* (MIM 182389), *SCN9A* (MIM 603415), *CACNA1A* (MIM 601011), *CACNA1C* (MIM 114205)) can cause developmental delay, intellectual disability, seizures, myotonia, ataxia, pain hypersensitivity and insensitivity, and cardiac arrhythmias. These conditions are far more common than CLIFAHDD syndrome, hundreds of different mutations have been reported to date, and for some conditions, significant structural-functional relationships have been described^35,36^. Caution is warranted, however, in trying to use these observations to make inferences about structural-functional relationships in *NALCN* because *NALCN* is not a paralog of these genes but instead diverged from them prior to the diversification of 4x6TM voltage-gated calcium and sodium channels in animals^37,38^.

In 2003, we reported that mutations in *TNNI2* and *TPM2* caused DA1 and hypothesized that distal arthrogryposis syndromes are in general caused by mutations in genes that encode components of the contractile apparatus of fast-twitch skeletal myofibers^1^. Over the past decade, we and others identified, via a candidate gene approach, mutations that cause DA conditions in four additional genes (i.e., *TNNT3*^2^, *MYH3*^3^, *MYH8*^39^, *MYBPC1*^40^ (MIM 160794)) that encode contractile proteins. However, mutations in these six genes collectively explain only ~60% of families with DA. In the past two years, exome sequencing has facilitated discovery of mutations in two genes, *PIEZO2* (MIM 613629), which cause distal arthrogryposis type 3 (Gordon syndrome; MIM 114300)^41^ and distal arthrogryposis type 5 (MIM 108145),^41,42^ and *ECEL1* (MIM 605896), which cause distal arthrogryposis type 5D (MIM 615065)^43,44^. In contrast to genes that encode contractile proteins, PIEZO2 is a mechanoreceptor and ECEL1 plays a role in the development of terminal neuronal branches at skeletal muscle endplates. These discoveries confirmed that a subset of DA syndromes is due to a mechanism(s) other than perturbation of the contractile apparatus of skeletal myofibers. The finding that mutations in *NALCN* underlie some persons diagnosed with an atypical DA, most often DA2A with neurological impairment, suggests that *NALCN* should be added to the expanding list of causal genes for DA conditions that do not encode contractile proteins. However, in each person with a *NALCN* mutation, the diagnosis of a DA should have been excluded because neurological abnormalities, including hypotonia and developmental delay, are explicit exclusion criteria^45^. In other words, CLIFAHDD is a newly delineated Mendelian condition and one of more than three hundred conditions characterized by arthrogryposis, but it is not a DA syndrome.

Descriptions of several children with clinical characteristics similar to CLIFAHDD syndrome have been reported previously. Specifically in the early 1990’s, four independent simplex families were described^46,47^ with “distal arthrogryposis, mental retardation, whistling face, and Pierre Robin sequence,” currently designated as Illum syndrome (MIM 208155). Three of these children died prematurely, including two who died at six months of age. In each instance, the facial characteristics, neurological findings, and natural history parallel those found in CLIFAHDD syndrome. Accordingly, we suspect that these children likely had mutations in *NALCN.* However, we note that Illum et al.^48^ reported a family with three affected siblings who had overlapping but distinctly different phenotypic features than are found in CLIFAHDD syndrome, including, most notably, calcium deposition in the skeletal muscle and central nervous system, that in combination suggest that Illum and CLIFAHDD syndromes are different conditions.

In summary, we used exome and targeted next-generation sequencing to identify *de novo* mutations in *NALCN* as the cause of a newly-delineated condition, CLIFAHDD syndrome, characterized by congenital contractures of the limbs and face, hypotonia, and developmental delay. The range of phenotypic variation in the persons with CLIFAHDD syndrome described herein is relatively narrow, but the congenital contractures were mild enough in some cases to have been overlooked, particularly in older children and young adults. Accordingly, it is possible that mutations in *NALCN* may also explain some cases of apparently isolated intellectual disability. Furthermore, the finding that the *de novo* mutations cluster in and around the pore-forming regions (i.e., S5 and S6 helices) suggests that the location and type of *NALCN* mutations may determine whether an individual develops IHPRF or CLIFAHDD syndrome.

## Supplemental Data

Supplemental Data includes four figures and a table.

## Acknowledgements

We thank the families for their participation and support; Christa Poel, Karynne Patterson, and Ian Glass for technical assistance; and J. David Spafford for helpful comments. Our work was supported in part by grants from the National Institutes of Health / National Human Genome Research Institute (1U54HG006493 to M.B., D.N., and J.S.; 1RC2HG005608 to M.B., D.N., and J.S.; 5R000HG004316 to H.K.T.), National Institute of Child Health and Human Development (1R01HD048895 to M.J.B. and 5K23HD057331 to A.E.B), the Life Sciences Discovery Fund (2065508 and 0905001), and the Washington Research Foundation.

## Web resources

The URLs for data presented herein are as follows:

Exome Variant Server (NHLBI Exome Sequencing Project ESP6500): http://evs.gs.washington.edu/EVS/

ExAC: http://exac.broadinstitute.org

Human Genome Variation: http://www.hgvs.org/mutnomen/

Online Mendelian Inheritance in Man (OMIM): http://www.omim.org/

SeattleSeq: http://snp.gs.washington.edu/

